# Large-scale deployment and establishment of *Wolbachia* into the *Aedes aegypti* population in Rio de Janeiro, Brazil

**DOI:** 10.1101/2021.04.29.441982

**Authors:** João Silveira Moledo Gesto, Sofia Pinto, Fernando Braga Stehling Dias, Julia Peixoto, Guilherme Costa, Simon Kutcher, Jacqui Montgomery, Benjamin R. Green, Katherine L. Anders, Peter A. Ryan, Cameron P. Simmons, Scott L. O’Neill, Luciano Andrade Moreira

## Abstract

Traditional methods of vector control have proven insufficient to reduce the alarming incidence of Dengue, Zika and chikungunya in endemic countries. The bacterium symbiont *Wolbachia* has emerged as an efficient pathogen-blocking and self-dispersing agent that reduces the vectorial potential of *Aedes aegypti* populations and potentially impairs arboviral disease transmission. In this work, we report the results of a large-scale *Wolbachia* intervention in Ilha do Governador, Rio de Janeiro, Brazil. *w*Mel-infected adults were released across residential areas between August 2017 and March 2020. Over 131 weeks, including release and post-release phases, we monitored the *w*Mel prevalence in field specimens, and analyzed introgression profiles of two assigned intervention areas, RJ1 and RJ2. Our results revealed that *w*Mel successfully invaded both areas, reaching overall infection rates of 50-70% in RJ1, and 30-60% in RJ2 by the end of the monitoring period. At the neighborhood-level, *w*Mel introgression was heterogeneous in both RJ1 and RJ2, with some profiles sustaining a consistent increase in infection rates and others failing to elicit the same. Correlation analysis revealed a weak overall association between RJ1 and RJ2 (*r* = 0.2849, *P* = 0.0236), and an association at a higher degree when comparing different deployment strategies, vehicle or backpack-assisted, within RJ1 (*r* = 0.4676, *P* < 0.0001) or RJ2 (*r* = 0.6263, *P* < 0.0001). The frequency *knockdown resistance* (*kdr*) alleles in *w*Mel-infected specimens from both areas was consistently high over this study. Altogether, these findings corroborate that *w*Mel can be successfully deployed at large-scale as part of vector control intervention strategies, and provide the basis for imminent disease impact studies in Southeastern Brazil.

## Introduction

Fighting the mosquito *Aedes aegypti* (= *Stegomyia aegypti*) sounds almost like a mantra for human populations living the tropics, whose lives are constantly threatened by diseases attributed to this species. Dengue (DENV), Zika (ZIKV) and chikungunya (CHIKV) viruses are among the many etiological agents transmitted by *Ae. aegypti*, highlighting its status as a major disease vector (Kraemer et al. 2015; WHO 2017). Global estimates of DENV alone point to around 400 million annual infections (Bhatt et al. 2013), distributed in over 128 countries (Brady et al. 2012). While the largest burden is in Asia (Bhatt et al. 2013), South American countries have long been hit by outbreaks and account for a considerable quota. In Brazil, notified cases of DENV sum up to 1.5 million annually, according to current surveillance reports (SVS 2019; 2021).

Without effective vaccines to tackle arboviral infections, public health authorities rely exclusively on vector control strategies (Thisyakorn and Thisyakorn 2014; Abdelnabi, Neyts, and Delang 2015; Lin et al. 2018). Management of breeding sites and deployment of chemical pesticides are the most common suppression methods, both with serious pitfalls. The former, usually performed by public agents and community members themselves, lacks precision and workforce, as suitable sites are vast in urban landscapes (Valença et al. 2013; Carvalho and Moreira 2017). In addition, *Ae. aegypti* egg loads are difficult to spot and remain viable for many months in nature (Rezende et al. 2008). As for the latter, natural selection of resistant variants has been the real issue (Maciel-de-Freitas et al. 2014; Melo Costa et al. 2020), downplaying the efficacy of current compounds and constantly pushing their replacement by new ones. Thus, innovative strategies tackling these issues and providing a more efficient, sustainable, control over arboviral infection are a welcome addition to traditional approaches in use.

One such strategy is the field deployment of *Wolbachia*-infected *Ae. aegypti. Wolbachia pipientis* is an obligatory intracellular bacterial endosymbiont, naturally present in around 40% of arthropods (Zug and Hammerstein 2012), which manipulates host reproductive biology to increase its inheritance rates (Werren, Baldo, and Clark 2008). When artificially introduced into *Ae. aegypti*, some *Wolbachia* strains such as *w*Mel or the virulent *w*MelPop were able to trigger cytoplasmic incompatibility (CI) in reciprocal crosses with wild specimens, and rapidly invade confined populations (Walker et al. 2011). In addition, and of particular importance to arboviral disease control, these newly established *Wolbachia*-mosquito associations led to pathogen interference (PI) phenotypes, possibly involving the modulation of immune system (Rancès et al. 2012) and metabolite pathways (i.e. intracellular cholesterol) (Caragata et al. 2014; Geoghegan et al. 2017). *Wolbachia*-harboring *Ae. aegypti* lines have shown refractoriness to infection by DENV, ZIKV, CHIKV and other medically relevant arboviruses (Moreira et al. 2009; Ferguson et al. 2015; Dutra et al. 2016; Aliota, Peinado, et al. 2016; Aliota, Walker, et al. 2016; Pereira et al. 2018; Carrington et al. 2018; Flores et al. 2020). Levels of refractoriness, nonetheless, seem to vary between strains, with a putative tradeoff with fitness costs (Joubert et al. 2016).

Supported by promising experimental data, *w*Mel-infected *Ae. aegypti* were used in pioneer field release trials in Northern Australia, promoting the bacterium spread and establishment into natural mosquito populations (A. A. Hoffmann et al. 2011; Ary A. Hoffmann et al. 2014). Importantly, *Wolbachia*’s high prevalence rates in the field, as well as intrinsic CI and PI, were sustained in the long-term, providing the necessary conditions to reduce dengue incidence in subsequent epidemiological assessments (O’Neill et al. 2019; Ryan et al. 2019). Corroborating the Australian findings, recent trials in Indonesia (Tantowijoyo et al. 2020) and Southeastern Brazil (Garcia et al. 2019; Gesto et al. 2020) have also reported the successful invasion and establishment of *w*Mel at some localities, with preliminary evidence of arboviral disease reduction (Indriani et al. 2020; Durovni et al. 2020; Pinto et al. 2021). In the particular context of Southeastern Brazil, trials have initially targeted small neighborhoods of Rio de Janeiro and the nearby city Niterói, following adult (Garcia et al. 2019) or egg deployment methods (Gesto et al. 2020). With high *w*Mel frequencies, and DENV and ZIKV refractoriness maintained intact over the post-release period (Gesto et al. 2020), additional areas of both cities could be considered for *Wolbachia* implementation.

In this study, we report the results of a large-scale field release of *w*Mel-infected *Ae. aegypti* in Rio de Janeiro, covering all the populated area of Ilha do Governador. We analyze the *Wolbachia* introgression profile, both from an overall and a more detailed neighborhood-specific perspective. To control for known operational risks we assess the *knockdown resistance* (*kdr*) profiling of colony and field specimens during our intervention. Lastly, we compare the outcomes of different adult deployment methods, ‘vehicle’ or ‘backpack’, and relate them to different urban and social contexts.

## Results and discussion

To evaluate the performance of a large-scale field deployment of *Wolbachia* in Brazil, we targeted the whole urban territory of Ilha do Governador, Rio de Janeiro. Being the largest island of Guanabara bay, with an estimated population of 211,018 and a total area of 40 km^2^, Ilha do Governador was an ideal starting point for testing expanded deployment interventions. First, because one of its neighborhoods, Tubiacanga, hosted a successful small trial in recent years (Garcia et al. 2019). And second, because islands are less prone to migration of wild mosquitoes from adjacent areas, which could affect the invasion dynamics. For logistical reasons, Ilha do Governador was divided into two great intervention areas, RJ1 and RJ2, each comprising a subset of neighborhoods (Figure 1). An additional layer was added by allocating sections to different deployment strategies: vehicle (V) or backpack-assisted (B). The former was the preferred choice, delivering speed and coverage, but was limited to areas with proper road organization, in which minivans could circulate and reach release sites. The latter was chosen for community settlements with informal housing and narrow passages, usually associated with favelas (i.e. slums). In this case, release sites could only be reached on foot.

**Figure 1.**
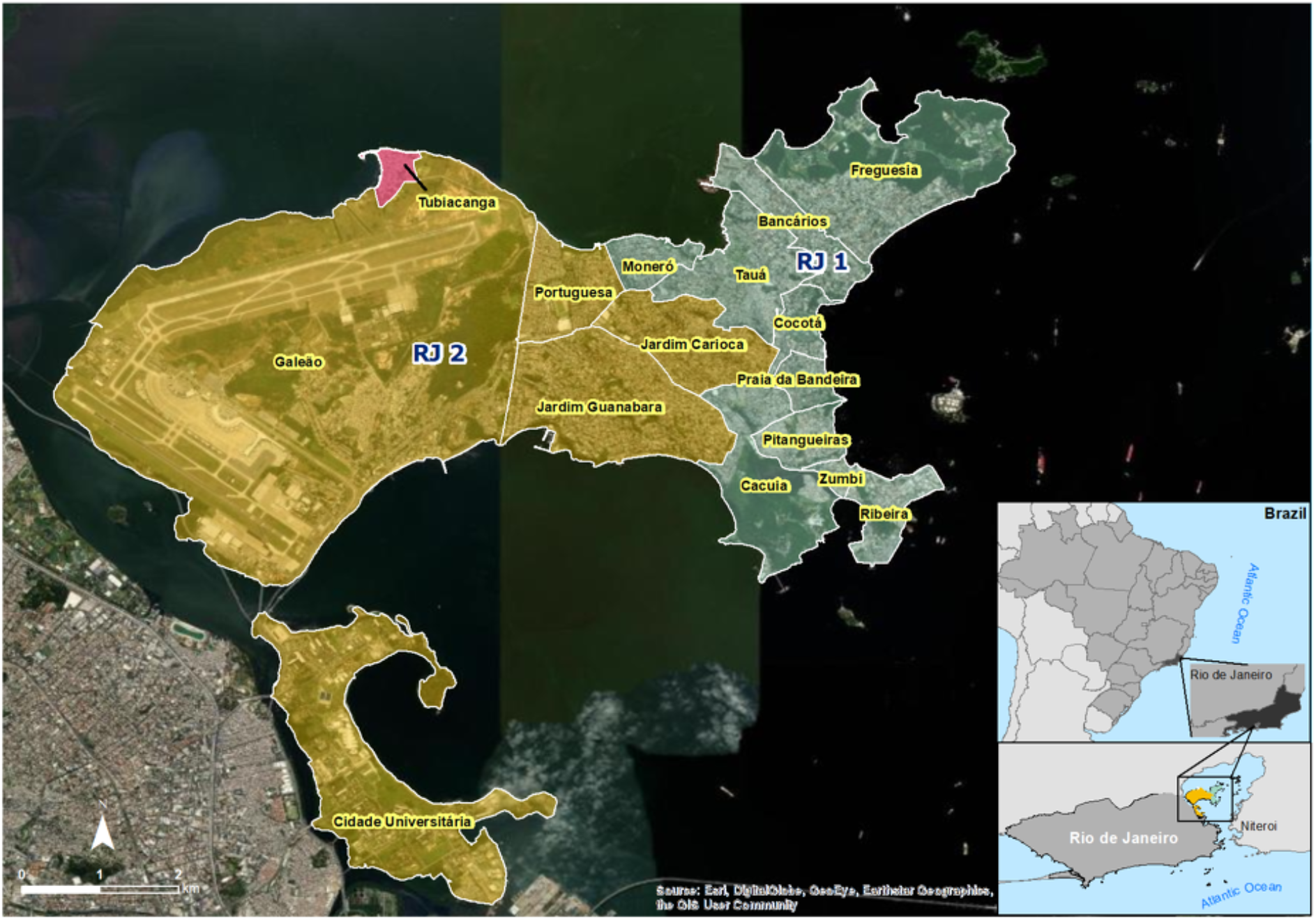
Map of Ilha do Governador intervention areas and neighborhoods. Satellite view of Ilha do Governador area, the largest island of Guanabara bay, in northern Rio de Janeiro (RJ). With an estimated total population of 211,018 in 40 km^2^, Ilha do Governador is divided into the following neighborhoods: Bancários, Cacuia, Cocotá, Freguesia, Monero, Pitangueiras, Praia da Bandeira, Ribeira, Tauá, Zumbi, Cidade Universitária, Galeão, Jardim Carioca, Jardim Guanabara, and Portuguesa. For *Wolbachia* release intervention, neighborhoods were grouped into two great areas, RJ1 (green) and RJ2 (yellow). Note that Cidade Universitária is actually located in an adjacent island, Ilha do Fundão, which is under the same public administration zone of Ilha do Governador and was therefore included as part of the RJ2 area. Also depicted is Tubiacanga (red), a small neighborhood which was targeted in a pioneer release trial.

*w*Mel-infected *Ae. aegypti* (*w*MelRio) (Garcia et al. 2019) were mass-released across RJ1 and RJ2, following specific schedules for each area (Figure 2, Figure S1, Supplementary Table S1). Mosquito deployments were carried in three rounds with ‘resting’ periods in-between, from August 2017 to March 2019 in RJ1, and from November 2017 to March 2019 in RJ2. To monitor *Wolbachia* presence in the field, BG-Sentinel traps were mounted in suitable households (Figure S2, Supplementary Table S2) and adult *Ae. aegypti* caught were tested weekly/fortnightly for *w*Mel infection. By analyzing the frequency of positive individuals (i.e. prevalence rates) over the 131 weeks spanning the entire release and post-release phases, the introgression of *w*Mel in RJ1 and RJ2, and across Ilha do Governador as a whole, were analysed (Figure 2).

**Figure 2.**
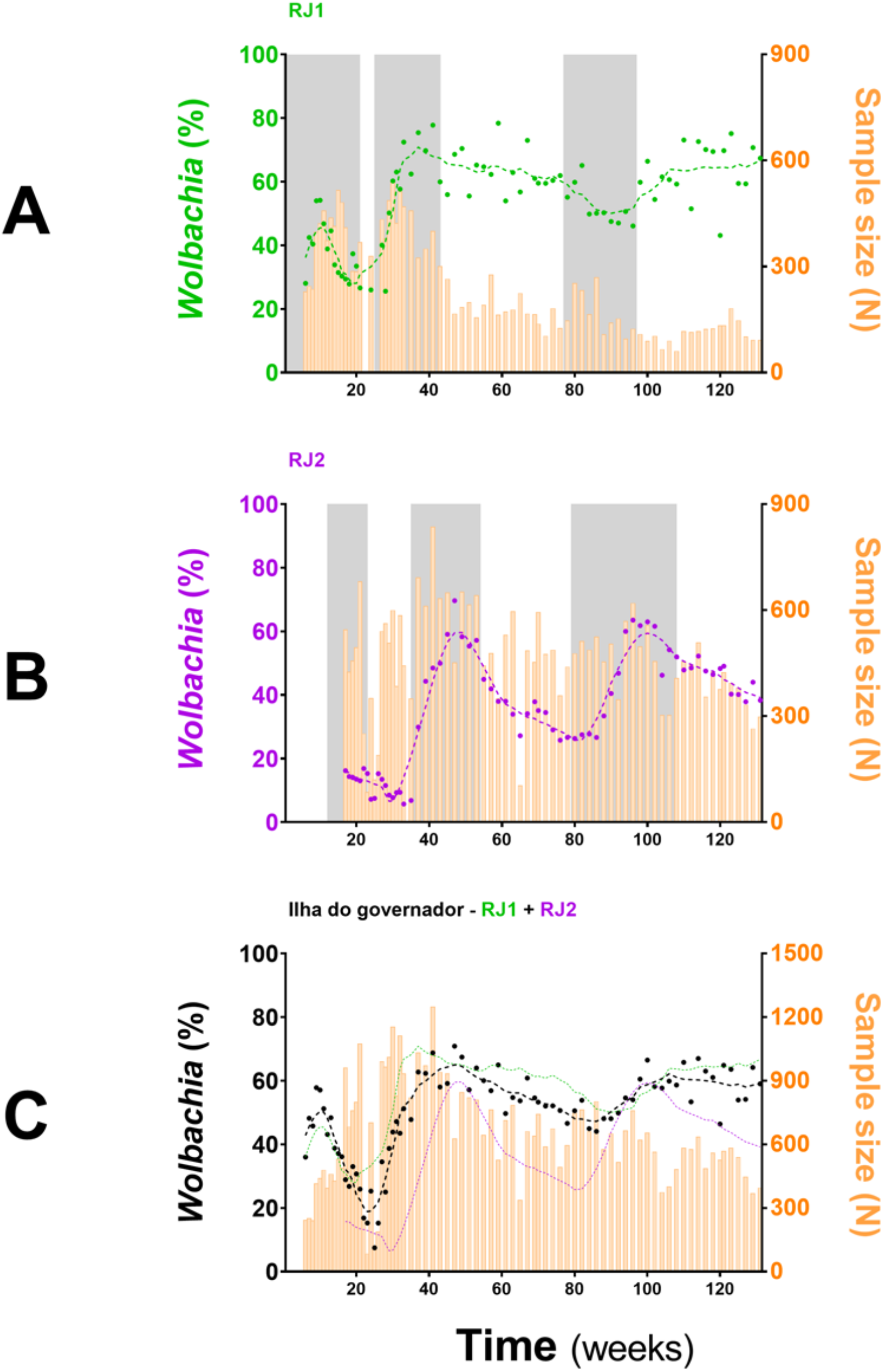
*Wolbachia*’s introgression into Ilha do Governador. Adult *w*Mel-harboring *Ae. aegypti* were mass-released in RJ1 and RJ2 areas, covering the entire territory of Ilha do Governador (Rio de Janeiro). Release intervention was carried out in three rounds (grey shading). Invasion profiles are depicted separately for (A) RJ1 and (B) RJ2, plus an aggregate of both for (C) an overall representation. Following the left Y-axis, *Wolbachia* prevalence indexes (%) are color coded and plotted as dots plus 2^nd^ degree, 7-neighbors, moving averages (dashed lines). Following the right Y-axis, sample sizes are plotted as histograms (orange bars). Time (weeks), since the beginning of adult releases (Week 1, August 2017) until recent days (Week 131, March 2020), is represented in the X-axis. Ticks are scaled for 20-week bins.

*<u>w</u>*Mel introgression in RJ1 was characterized by a steep increase in prevalence rates over the first two release rounds, peaking at 60-80%, and a subsequent and self-sustaining frequency of 50-70%, until the end of monitoring (Figure 2A). At the neighborhood level (Figure 3) *<u>w</u>*Mel introgression was heterogenous in RJ1, suggesting that invasion dynamics were not consistent across the intervention area. In some of the neighborhoods where releases were vehicle-assisted, such as Bancários, Freguesia, Monero, Tauá, Cacuia and Praia da Bandeira the introgression profiles showed consistent increases in *w*Mel prevalence over time, resulting in high prevalence rates (>80%) by March 2020. Others, such as Pitangueiras, Cocotá and Ribeira, had more heterogenous profiles, with alternating mid-high (60-80%) and low (<50%) *w*Mel frequencies over time, with non-consistent trends by the end of March 2020. The neighborhood of Zumbi reached moderately high *w*Mel prevalence (60-70%) following the second round of releases, but monitoring was then suspended in March 2019 due to very low mosquito abundance, precluding further observation. The areas with backpack-assisted releases, aggregated and analyzed as a single unit named RJ1.B, showed a slow and consistent rise in *w*Mel prevalence towards high levels (>80%), suggesting successful wMel introgression (Figure 3).

**Figure 3.**
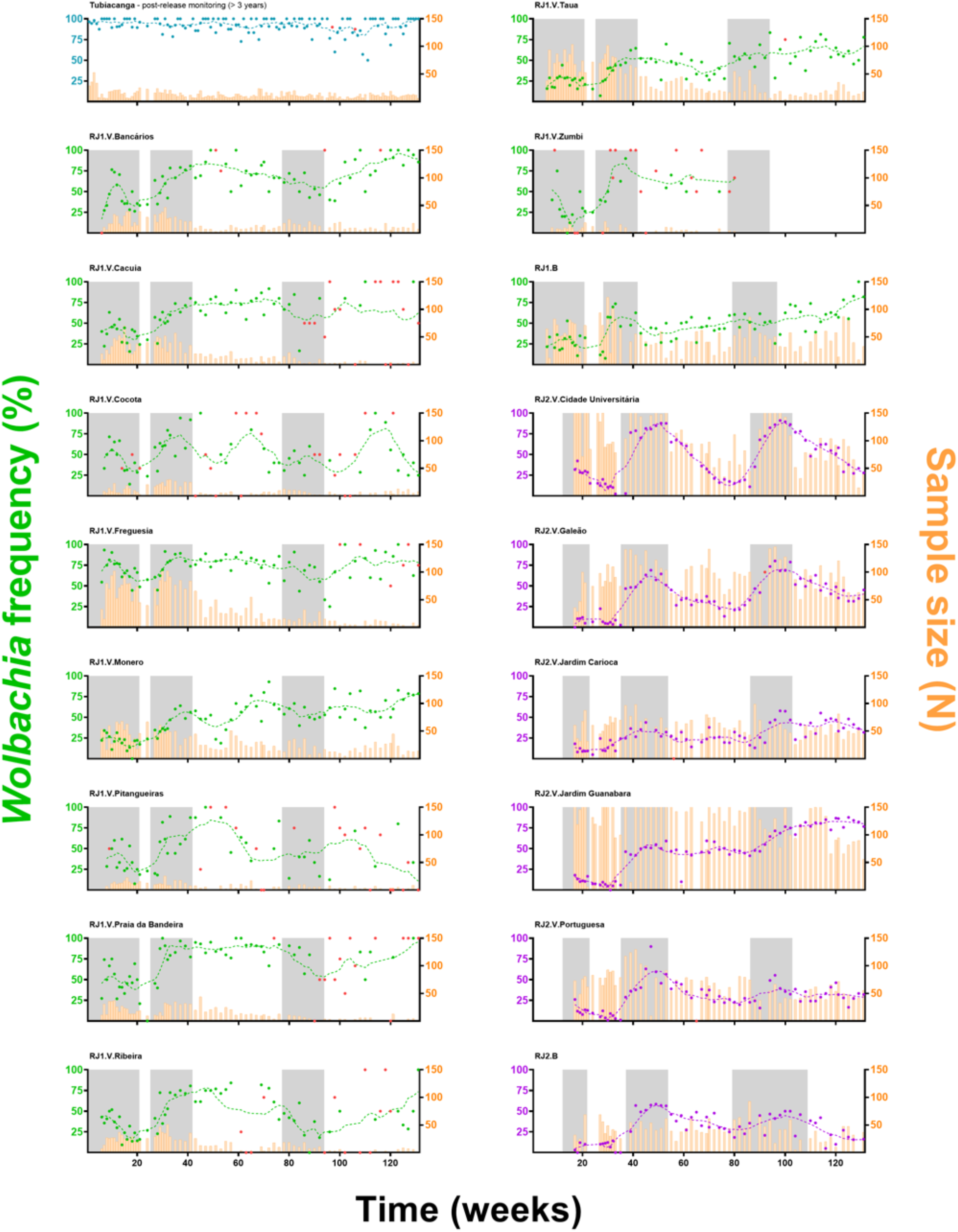
*Wolbachia*’s invasion profiles in individual neighborhoods. Adult *w*Mel-harboring *Ae. aegypti* were released (grey shading) across all neighborhoods of Ilha do Governador. Individual invasion profiles are depicted, with ‘RJ1’ (green) or ‘RJ2’ (purple) coding for the intervention area, and ‘V’ or ‘B’ for vehicle or backpack-assisted releases, respectively. *Wolbachia* prevalence indexes (%) are color coded and represented by dots plus 2^nd^ degree, 7-neighbors, moving averages (dashed lines), following the left Y-axis. Sample sizes are plotted as histograms (orange), following the right Y-axis. Prevalence indexes from small-sized samples (N<5) are marked in red. The X-axis represents time (weeks), since the beginning of adult releases (Week1, August 2017) until recent days of field monitoring (Week 131, March 2020), with ticks scaled accordingly to represent 20-week bins. Post-release *Wolbachia* prevalence in Tubiacanga (blue), a previous intervention site, is shown as a standard for long-term field establishment.

In RJ2, the overall introgression profile was characterized by oscillating *w*Mel frequencies (30-60%), with prevalence rates increasing over the second and third release rounds but not self-sustaining afterwards (Figure 2B). Once again, individual neighborhood results indicate a complex, spatially variable picture of wMel introgression (Figure 3). Here, in vehicle-assisted release areas, Jardim Guanabara was the best performing, with a classical invasion trend stabilizing at high prevalence rates (∼80%). Jardim Carioca and Portuguesa, on the other hand, were less successful and had persistently low frequencies (30-40%). Unlike the two categories above, Galeão and Cidade Universitária had mid-level wMel frequencies (30-60%), similar to the overall RJ2 profile. These two neighborhoods account for most of the territory enclosed in RJ2, with sparse building blocks and a peculiar, mostly non-resident human occupation. Galeão hosts the city’s international airport, and Cidade Universitária, as the name suggests, hosts the federal university campus. In backpack-assisted release areas, RJ2.B, prevalence rates also increased during second and third release rounds, but soon after declined to low levels and, at the time of last monitoring in March 2020, did not yet demonstrate evidence of *w*Mel introgression.

Despite intrinsic differences in their overall profiles, RJ1 and RJ2 are still weakly associated in Spearman’s correlation analysis (*r* = 0.2849, *P* = 0.0236) (Figure 4A), suggesting that factors underlying invasion are shared at some level between intervention areas. Hence, RJ1 and RJ2 data were aggregated into a single profile reflecting the overall panorama of *Wolbachia*’s invasion in Ilha do Governador (Figure 2C). With prevalence rates ranging from 55 to 65% by the end of field monitoring, this panorama suggests that *w*Mel introgression is still an on-going process in Ilha do Governador. This representation, however, must be understood as an oversimplified indicative of its invasion dynamics, hiding an underlying complexity at the neighborhood (or neighborhood section) level.

**Figure 4.**
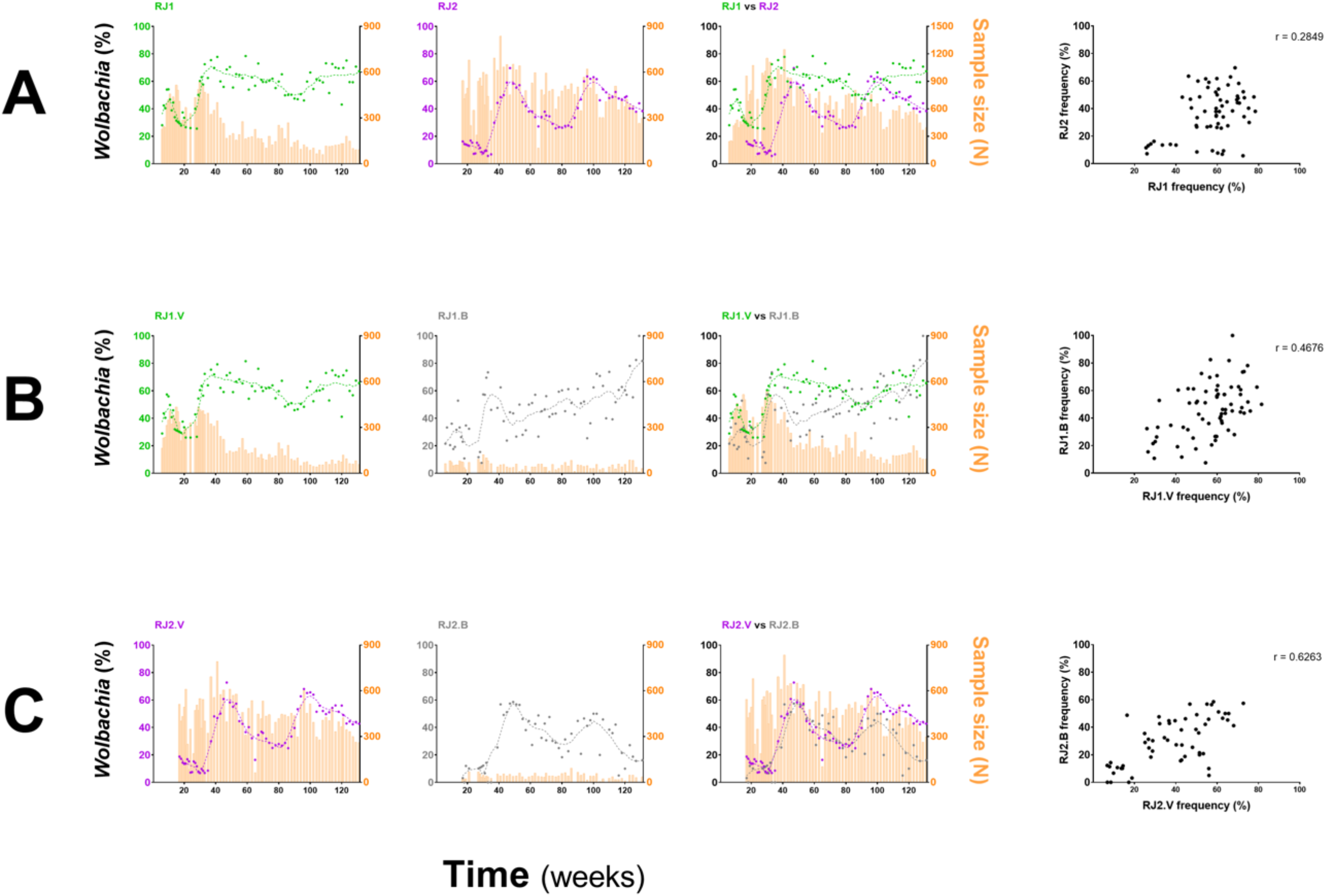
Comparison of invasion profiles between intervention areas and deployment strategies. *w*Mel frequencies for different intervention areas, or deployment strategies, were represented individually and overlayed, and compared by Spearman’s correlation analyses. A) RJ1 vs RJ2; B) RJ1.V vs RJ1.B; C) RJ2.V vs RJ2.B. The degree of association between frequency datasets is indicated by the *r* coefficient, at the top right of the correlation graphs.

A similar analysis was undertaken to compare vehicle and backpack-assisted sections, within RJ1 (Figure 4B) and RJ2 contexts (Figure 4C). Spearman’s correlation indicates a moderate association between RJ1.V and RJ.B (*r* = 0.4676, *P* < 0.0001), and a strong association between RJ2.V and RJ2.B (*r* = 0.6263, *P* < 0.0001), suggesting that the outcomes of both deployment strategies covary within the same region. Nonetheless, the efficiency of each strategy, which ultimately translates to weekly prevalence rates and invasion trends, was variable between intervention areas and possibly affected by non-controlled events. In backpack-assisted sections, RJ1.B and RJ2.B, release intervention was often impaired by violent drug-related conflicts. During the third round, RJ1.B had 5 weeks of interruption due to this reason alone, and RJ2.B had 6, extending the release period to 17 or 28 weeks, respectively (Supplementary Table S1). Interestingly, should interruptions of this kind influence invasion dynamics, then RJ2.B was certainly more affected, as revealed by our failed attempt to stably introgress *w*Mel by the end of this study period (i.e. week 131).

We previously deployed *w*Mel in the small community of Tubiacanga on the Ilha do Governador along 2014 and 2015. Here, *w*Mel initially failed to establish because of mismatched genetic backgrounds between the release mosquito strain and the resident wild-type population, raising particular attention to insecticide resistance-related traits (Garcia et al. 2019). It was only after repetitive rounds of backcrossing, with introgression of wild allelic variants, that the *w*Mel-infected line was enough fit to promote a successful invasion. With this in mind, we monitored the genetics of pyrethroid resistance by screening for mutations in the voltage-gated sodium channel (*Na_v_*), known as *kdr* (knockdown resistance), in field caught *Wolbachia*^+^ samples from RJ1 and RJ2 (Figure 5). As we could observe, the allelic profiling of samples from both intervention areas revealed the predominance of resistant variants, *Na_v_R1*(1016Val^+^, 1534Cys^kdr^) and *Na_v_R2* (1016Ile^kdr^, 1534Cys^kdr^), and shortage of the susceptible one, *Na_v_S* (1016Val^+^, 1534Phe^+^), corroborating the findings of a nation-wide survey (Melo Costa et al. 2020). Over the monitoring period, this profile experienced little variation within and between areas, indicating the long-term maintenance of *kdr* mutations in *w*Mel-infected field samples, and highlighting its adaptive role in pyrethroid-infested environments. Moreover, it rules out the possibility that the differences observed in invasion trends along this trial could be influenced by *kdr* frequencies.

**Figure 5.**
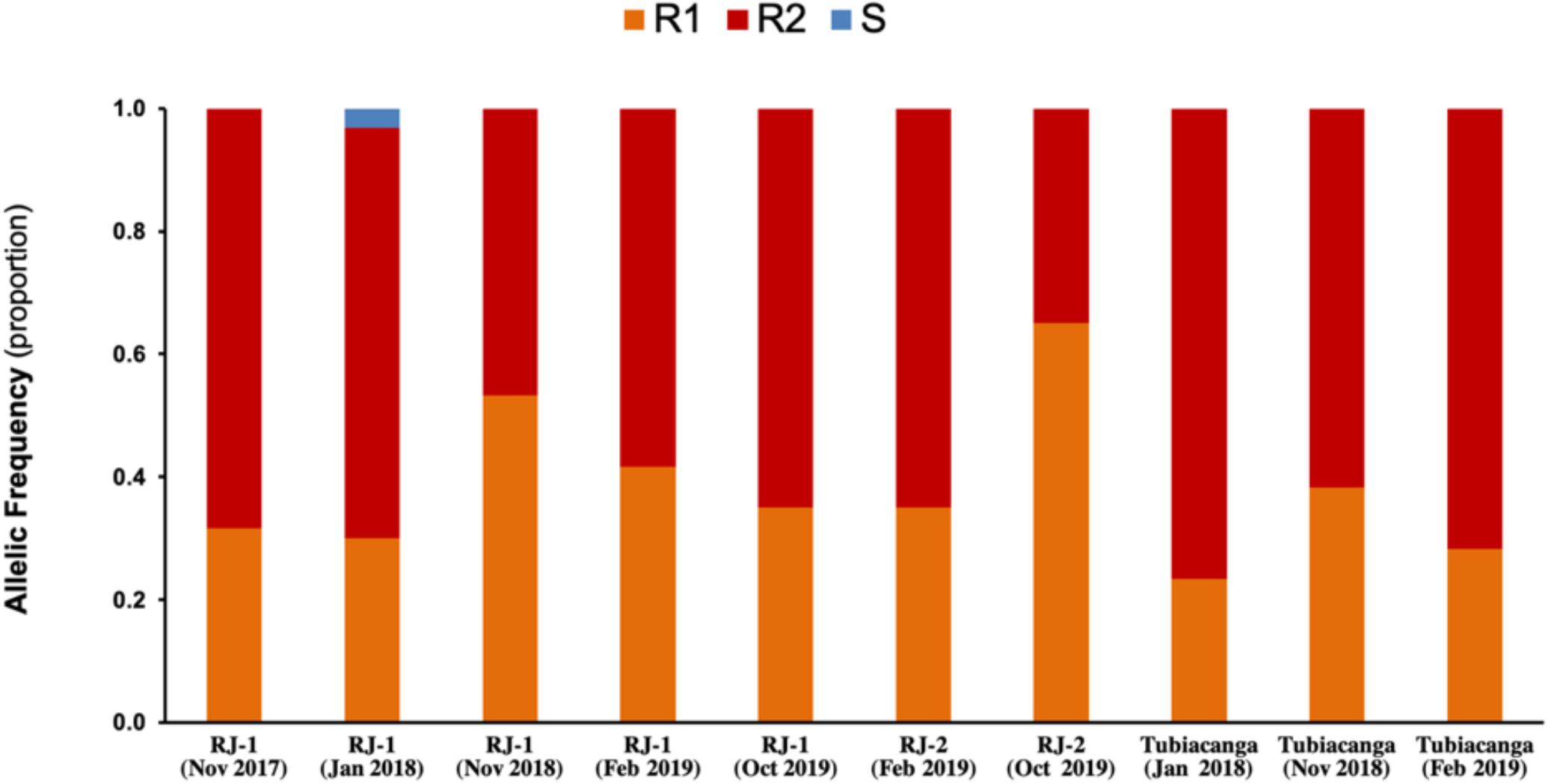
Genetic monitoring of insecticide resistance in intervention areas. Knockdown resistance (*kdr*) allelic variants were monitored in field caught mosquitoes (*Wolbachia*^+^) over the release and post-release interventions in RJ1 and RJ2. Data represent the proportion of alleles linked to susceptibility, S, or resistance to insecticides, R1 and R2, in each sample (n=30). Mosquitoes from Tubicanga, home to a previous successful trial, were also included for comparison. S (*Na_v_S*): 1016Val^+^; R1 (*Na_v_R1*): 1016Val^+^, 1534Cys^kdr^; R2 (*Na_v_R2*): 1016Ile^kdr^, 1534Cys^kdr^.

To drive a successful invasion, *Wolbachia* must interact with bacterium-free *Ae. aegypti* populations and underlying factors that influence its maintenance and density in the natural habitat (Hancock et al. 2016; Schmidt et al. 2017). Here, especial attention should be given to quiescent egg loads, which are known to remain viable for many months (up to over a year) in the habitat, waiting for favorable conditions to resume. With a reduced resistance to desiccation, *w*Mel-infected eggs are critically impacted by climate and have a significant decay in viability in periods over 40 days (Farnesi et al. 2019). Although it is not clear how much it costs to invasion profiles, it is still an underlying factor to consider when analyzing different contexts. From this perspective, human settlements with fewer inhabitants and/or better management of breeding sites, aided by community participation in vector control surveillance, could be prone to lower *Ae. aegypti* densities and faster, more efficient, *Wolbachia* invasion. In contrast, crowded human settlements and undermined control of breeding sites tend to promote higher *Ae. aegypti* densities and slower, less efficient, invasion dynamics.

Even though some individual neighborhoods of Ilha do Governador failed to elicit invasion trends, it is possible that this scenario reverts on its own in the future. Here, migration from adjacent neighborhoods (Schmidt et al. 2017), with higher prevalence rates, may play an important contribution and act synergistically with *w*Mel self-driving ability, as expressed by the CI mechanism. In other words, *Wolbachia* hotspots like Bancários, Freguesia, Monero, Praia da Bandeira, Jardim Guanabara and Tubiacanga could serve as autonomous centers to deliver migrants to less prevalent neighborhoods, helping them to achieve high and sustainable rates in the future, and providing a more uniform establishment into Ilha do Governador. This effect, however, can only be verified after a continued long-term monitoring of field populations, whose data may also indicate the necessity to apply topic release boosts at those neighborhoods with persistent low rates. These considerations are part of challenging large-scale release interventions, which are still incipient here in Brazil and in other parts of the world (Schmidt et al. 2017; Ryan et al. 2019; Tantowijoyo et al. 2020). As a result of accumulating data from current trials, we shall better understand the factors underlying invasion dynamics and optimize future strategies.

Altogether, our results ratify that *w*Mel field release is adaptable to large-scale, using coordinated efforts to impact densely populated areas. With continuous improvement of rearing and release technology, it could be amplified to cover city-wide territories in short time. As preliminary disease impact studies suggest (Durovni et al. 2020; Pinto et al. 2021), one could foresee a significant reduction in the incidence of dengue and other arboviral diseases in Rio de Janeiro and nearby Niteroi, fulfilling the main goal of current trials and cementing *Wolbachia* as an efficient and sustainable solution vector control in Brazil.

## Methods

### Mosquito husbandry

To generate *w*MelRio, a precursor Australian line harboring the *w*Mel strain of *Wolbachia* (Walker et al. 2011) was backcrossed for eight generations to a natural *Ae. aegypti* population from Rio de Janeiro (Dutra et al. 2015). To achieve high genetic background homogenization, additional crosses followed by *knockdown resistance* (*kdr*) screening were performed, and individuals whose *kdr* profiling resembled that of the natural population were positively selected (Garcia et al. 2019). To prevent drift and selection of new variants in our facilities, and keep *w*MelRio in resonance with the natural background, our colony was refreshed every five generations with 10% wild-caught males.

*w*MelRio eggs were hatched in degasified water with 0.08% Tetramin^®^ (Tetra GmbH, Herrenteich, Germany). After 5 h incubation at room temperature, hatching rates were calculated and first instar (L1) larvae were transferred to mass-rearing trays containing 20-30k individuals each. Larval development (L1 to L4) was promoted at 28 °C, in dechlorinated water supplemented with liquid diet (3.7% fish meal, 2.6% liver powder and 1.1% brewer’s yeast). On the sixth day, with pupae formation reaching levels up to 50-70%, immatures were collected and sent to either colony renewal or mass-release pipelines (see details for the latter in ‘Adult releases’).

For colony renewal, immatures were split in groups of approximately 2,000 individuals and placed inside BugDorm^®^ cages (MegaView Science Co Ltd, Taiwan). Adult emergence and husbandry occurred at 25 °C, with 10% sucrose solution *ad libitum*. Females were fed human blood (from donation centers; more details in ‘ethical regulations’) every 2-3 days, through Hemotek^®^ artificial feeders (Hemotek Ltd, UK). Here, biosafety and ethical guidelines were followed to prevent arboviral contamination of our colony and mass-release batches, with all blood samples negatively scored for DENV, ZIKV, CHIKV, MAYV and YFV by multiplex qPCR (Dutra et al. 2016; Pereira et al. 2018). For egg-collection, dampened filter papers (i.e. half-immersed in water) were placed inside the cages for 2-3 days, before being removed and gradually dried at room temperature. Egg strips (a.k.a. ovistrips) were stored at room temperature until further use, either for colony maintenance or field release. Egg strips stored for more than 40 days were discarded due to a decay in overall quality (Farnesi et al. 2019).

### Adult releases

For the mass-release of *Wolbachia*-harboring *Ae. aegypti*, batches of approximately 150 late-stage immatures were transferred to release tubes: custom-made acrylic pipes closed at both ends with a fine mesh, allowing both liquid and air flow during the final developmental stages. Following adult emergence, release tubes were counted, quality assessed and designated to ‘backpack’ or ‘vehicle’ deployment.

For ‘vehicle’ deployment, release tubes were stacked into mini vans at dawn before departing to trips covering a fraction of the release sites in Ilha do Governador. Each van followed a strict routine, leaving the mass-production facility at scheduled times, and with the driver and the release agent fully aware of the map, traffic and possible turnarounds. When the van hit the approximate location of the sites, the agent would extend his/her arms outside the window and gently remove the mesh to free the adults kept inside the tube. Once completed, the van would proceed to the following site to repeat the procedure.

For ‘backpack’ deployment, release tubes were stacked inside backpacks before departing to areas with restricted access, either because of irregular housing and narrow alleys, or because of drug-related episodes of violence. In these areas, deployment was carried out on foot by public health agents, working in partnership with both the WMP staff and community leaders. As usual, before starting a trip, agents were given maps and routes to cover the release sites, and asked to report their activity and any obstacle that might arise by the end of the day.

The number and spatial distribution of release sites (Figure S1, Supplementary Table S1) was strategically defined so as promote an efficient spread of *w*Mel-harboring individuals into each neighborhood. Release sites were geotagged and integrated to ©OpenStreetMap source data with ArcGIS version 10.4 (Esri, Redlands, CA, USA), allowing the planning of daily routes and a better control and management over the whole release intervention. Schedules (Supplementary Table S1) varied according to the area, RJ1 or RJ2, and deployment method, ‘vehicle’ or ‘backpack’, being revisited after each round based on the status of *Wolbachia* frequency in the field. Additional rounds were applied in order boost the frequency levels and promote an efficient invasion.

### Ethical regulations

Regulatory approval for the field release of *Wolbachia*-harboring *Ae. aegypti* was obtained from the National Research Ethics Committee (CONEP, CAAE 02524513.0.1001.0008), following a common agreement of governmental agencies (IBAMA, Ministry of Environment; ANVISA, Ministry of Health; and MAPA, Ministry of Agriculture, Livestock and Supply) and the former sanction of the special temporary registry (RET, 25351.392108/2013-96). Community acceptance was evaluated by social engagement activities and a fill out questionnaire, with all neighborhoods recording > 70% household support. Written informed consents were acquired from those hosting BG-sentinel traps, who were offered financial aids to cover electricity costs.

Additional regulatory approval (CONEP, CAAE 59175616.2.0000.0008) was required to feed the adult female mosquitoes with human blood, a necessary step for the maintenance of *w*MelRio colony and mass production of eggs. We only used blood which would have been discarded by not attending quality assurance policies (e.g. blood bags with insufficient volume) of donation centers: Hospital Pedro Ernesto (Universidade Estadual do Rio de Janeiro) and Hospital Antonio Pedro (Universidade Federal Fluminense). All blood samples complied with Brazilian Government guidelines for routine screening, having no information on donor’s identity, sex, age and any clinical condition, as well as testing negative for Hepatitis B, Hepatitis C, Chagas disease, syphilis, HIV and HTLV.

### Field population monitoring and *Wolbachia* diagnosis

BG-Sentinel traps (Biogents AG, Regensburg, Germany) were spread across all neighborhoods of Ilha do Governador (RJ) to monitor the *Wolbachia* frequency in the field (Supplementary Figure S2, Supplementary Table S2). Monitoring sites covered an area of approximately 250 m^2^ each, and were selected among suitable households who formally accepted hosting a trap. For an optimal control over the monitoring area and map creation, sites were geotagged and overlayed with ^©^OpenStreetMap source data using ArcGIS version 10.4 (Esri, Redlands, CA, USA). Overtime, reallocation of sites was often necessary and occurred when households quit hosting the trap, or in cases of equipment misuse or failure. Staff agents checked each working trap weekly, bringing the catch bags (perforated envelopes positioned inside the BG-Sentinels to trap insects) to our facilities for species identification and *Wolbachia* screening.

*Ae. aegypti* samples were individually screened for *Wolbachia* by qPCR or LAMP. In short, whole-bodies were homogenized in Squash Buffer (10 mM Tris-Cl, 1 mM EDTA, 25 mM NaCl, pH 8.2) supplemented with Proteinase K (250 ug/ml). DNA extraction was carried out by incubating the homogenates at 56 °C for 5 min, followed by 98 °C for 15 min to stop the proteinase activity. qPCR reactions were performed with LightCycler® 480 Probes Master (Roche), using specific primers and probes to amplify *Wolbachia pipientis* WD0513 and *Ae. aegypti rps17* genes (Supplementary Table S3). Temperature cycling conditions were set on a LightCycler® 480 Instrument II (Roche), using the following parameters: 95 °C for 10 min (initial denaturation), and 40 cycles of 95 °C for 15 s and 60 °C for 30 s (single acquisition). LAMP (*Loop-Mediated Isothermal Amplification*) reactions were performed with WarmStart® Colorimetric LAMP 2X Master Mix (DNA & RNA) (New England Biolabs) and an alternative set of primers (Supplementary Table S3), as described elsewhere (Gonçalves et al. 2019). Isothermal amplification was carried out at 65 °C for 30 min on a T-100 Thermocycler (Bio-Rad), according with manufacturer conditions. Both qPCR and LAMP reactions were performed in 96-well plates. Specimens with and without *Wolbachia* were used as positive and negative controls, respectively.

### *kdr* genotyping

Adult *Ae. aegypti* were genotyped by qPCR to detect single nucleotide polymorphisms (SNPs) at the 1016 (Val^+^ or Ile^kdr^) and 1534 (Phe^+^ or Cys^kdr^) positions of the voltage gated sodium channel gene (*Na_v_*), as previously reported (Macoris et al. 2018; Hayd et al. 2020). Amplification reaction was performed with LightCycler 480 Probes Master mix (Roche), 10ng of individual genomic DNA, and a set of primers and probes to detect *kdr* alleles (Supplementary Table S3) customized by Thermo Fisher Inc. under ID codes: AHS1DL6 (Val^+^1016Ile^kdr^) and AHUADFA (Phe^+^1534^Cys^). Thermal cycling was carried out on a Light Cycler 480 Instrument II (Roche), set to the following conditions: 95 °C for 10 min (initial denaturation), and *N* cycles of 95 °C for 15 s and 60 °C for 30 s (single acquisition). *N* was set to 30, for amplifying Val^+^1016Ile^kdr^, or to 40, for Phe^+^1534Cys^kdr^. For each collection date, 30 samples were individually genotyped. Rockefeller colony specimens (kindly provided by Dr. Ademir de Jesus Martins Júnior, IOC, Fiocruz), harboring susceptible (*Na_v_S*) or resistant variants (*Na_v_R1* and *Na_v_R2*), were used as positive controls.

### Statistical Analyses

All statistical analyzes were performed in Graphpad Prism 8 (Graphpad Software, Inc). *Wolbachia* frequency time-series were smoothed using a moving average of 7-neighbors, 2^nd^ order polynomials. Spearman correlation *r* coefficient was used to compare invasion trends between great intervention areas, RJ1 and RJ2, as well as the deployment strategies applied within each, ‘vehicle’ or ‘backpack’. For all statistical inferences, α was set to 0.05.

## Data Availability Statement

All relevant data generated or analyzed during this study are included in this manuscript (and its Supplementary Information file).

## Ethics Statement

This study and all the underlying work regarding the rearing and field-release of *Wolbachia*-harboring *Ae. aegypti* was approved by the National Research Ethics Committee (CONEP, CAAE 02524513.0.1001.0008), following a common agreement of governmental agencies (IBAMA, Ministry of Environment; ANVISA, Ministry of Health; and MAPA, Ministry of Agriculture, Livestock and Supply) and the former sanction of the special temporary registry (RET, 25351.392108/2013-96).

## Author contributions

J.S.M.G., S.P., F.B.S.D., J.P., G.C. and L.A.M. conceived the study. J.S.M.G., S.P., F.B.S.D., J.P., G.C., S.K., J.M., B.R.G., P.A.R., C.P.S., K.L.A., S.L.O. and L.A.M. performed the investigation, data curation and analysis. L.A.M. managed the project supervision, validation and funding. J.S.M.G., S.P. and L.A.M. drafted the manuscript, and all authors reviewed and approved its final version.

## Funding

This work was funded by the Brazilian Ministry of Health, and a grant to Monash University from the Bill and Melinda Gates Foundation. The funders had no role in study design, data collection and analysis, decision to publish, or preparation of the manuscript.

## Conflict of Interest

The authors have declared that no conflict of interests exists.

## Acknowledgements

We acknowledge all team members of the World Mosquito Program, past and present, who were involved and genuinely believed in the *Wolbachia* method as solution to attenuate the burden of arboviral diseases in densely populated areas of Rio de Janeiro. The successful outcome of this study, as well as its future legacy, was only achieved because of their dedication and team-working. We thank Flavia Teixeira, for managing ethical and regulatory approvals and compliance, and Roberto Peres and Catia Cabral, for supervising the mass-rearing, field release and *Wolbachia* monitoring. Special thanks to WMP Global for overall advice and tech transfer underlying large-scale deployment. We are also very grateful to the public agents of the Health Municipality of Rio de Janeiro, for the providing essential field assistance, and to community members of Ilha do Governador, for actively supporting and engaging in our activities.

## Notes

### Competing Interest Statement

The authors have declared no competing interest.

