## Supplementary Information for "Large-scale deployment and establishment of *Wolbachia* into the *Aedes aegypti* population in Rio de Janeiro, Brazil"

**Supplementary Figure S1. Spatial distribution of adult-release sites.** Approximate locations of release sites are depicted separately for each neighborhood of Ilha do Governador, and according to the deployment strategy: vehicle (blue dots) or backpack (red dots). A) RJ1.Bancários, B) RJ1.Cacuia, C) RJ1.Cocotá, D) RJ1.Freguesia, E) RJ1.Monero, F) RJ1.Pitangueiras, G) RJ1.Praia da Bandeira, H) RJ1.Ribeira, I) RJ1.Tauá, J) RJ1.Zumbi, K) RJ2.Cidade Universitária, L) RJ2.Galeão, M) RJ2.Jardim Carioca, N) RJ2.Jardim Guanabara, O) RJ2.Portuguesa. Urban perimeters enclosing release sites (black lines) filter out inhabited areas with low-abundance *Ae. aegypti*. Geotagged release site maps were generated using ArcGIS 10.4 (Esri, Redlands, CA, USA) and ©OpenStreetMap source data. Scale is represented at the bottom left corner.

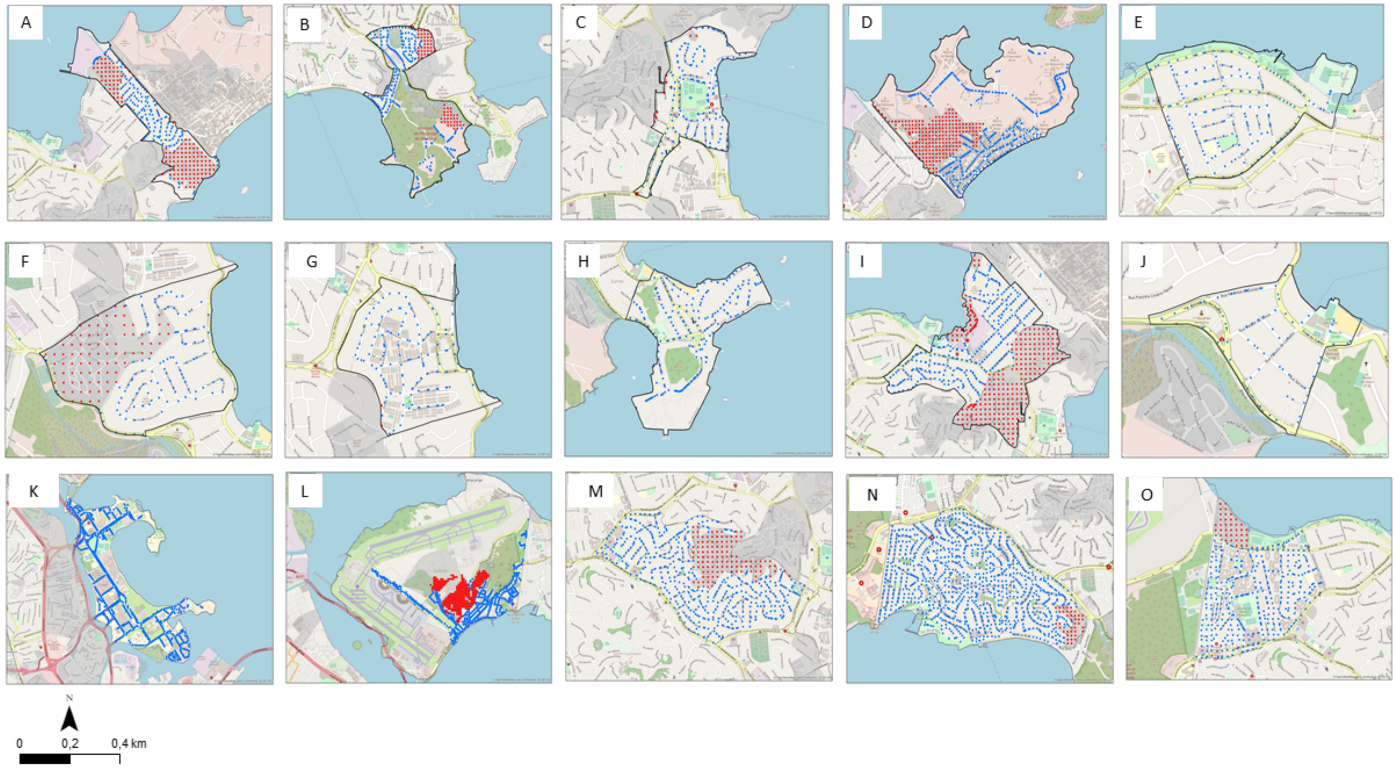

**Supplementary Figure S2. Spatial distribution of monitoring sites.** Geotagged locations of BG-sentinel traps set in household partners are depicted separately for each neighborhood of Ilha do Governador, according to deployment strategy used: vehicle (black dots) or backpack (purple dots). A) RJ1.Bancários, B) RJ1.Cacuia, C) RJ1.Cocotá, D) RJ1.Freguesia, E) RJ1.Monero, F) RJ1.Pitangueiras, G) RJ1.Praia da Bandeira, H) RJ1.Ribeira, I) RJ1.Tauá, J) RJ1.Zumbi, K) RJ2.Cidade Universitária, L) RJ2.Galeão, M) RJ2.Jardim Carioca, N) RJ2.Jardim Guanabara, O) RJ2.Portuguesa. Urban perimeters (black lines) filter out non-inhabited areas with low-abundance *Ae. aegypti*. Trapped specimens were assessed for *Wolbachia* frequency status. Maps were created with ArcGIS 10.4 (Esri, Redlands, CA, USA) and ©OpenStreetMap source data. Scale is represented at the bottom left corner.

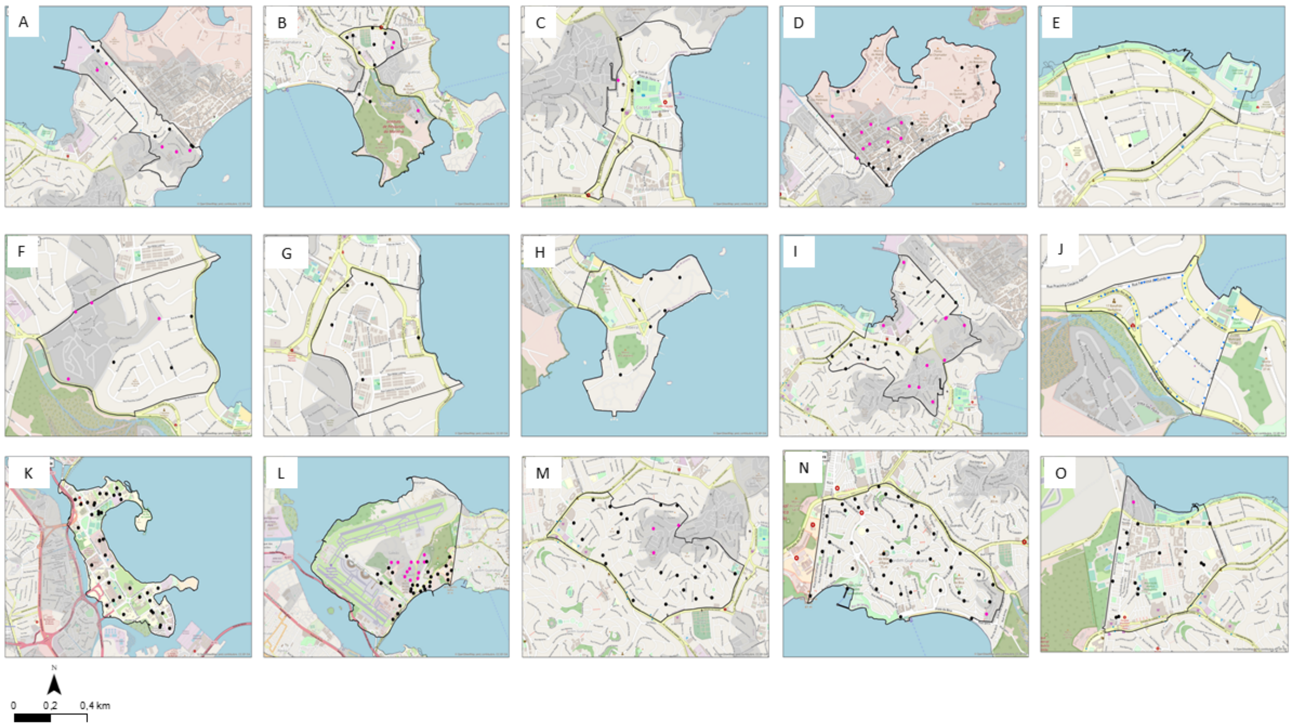

**Supplementary Table S1. Adult-release schedules and dosages across intervention areas.** Schedules and numbers of released mosquitoes per intervention area, RJ1 or RJ2, and deployment strategy, vehicle (V) or backpack-assisted (B).

| Area | Begin | End | Weeks | Events[a] | Sites/ event[b] (± SD) | Released/ event[c] (± SD) | Released/ event/ site[d] (± SD) | Released/ event/ area[e] (± SD) | Released (Total) |
|---|---|---|---|---|---|---|---|---|---|
| **1st Round** | | | | | | | | | |
| RJ1.B | 29/08/2018 | 01/02/2018 | 20 | 10 | 871 (± 0) | 100,601 (± 42,932) | 118 (± 45) | 48,836 (± 20,841) | 1,006,012 |
| RJ1.V | 29/08/2017 | 02/01/2018 | 20 | 14 | 1,668 (± 642) | 228,159 (± 93,244) | 137 (± 14) | 36,623 (± 14,967) | 3,194,229 |
| RJ2.B | 21/11/2017 | 16/01/2018 | 9 | 8 | 763 (± 0) | 77,462 (± 44,269) | 101 (± 58) | 56,132 (± 32,079) | 619,699 |
| RJ2.V | 23/11/2017 | 25/01/2018 | 10 | 9 | 4,657 (± 1,348) | 419,655 (± 329,660) | 91 (± 60) | 27,215 (± 21,379) | 3,776,895 |
| **2nd Round** | | | | | | | | | |
| RJ1.B | 13/03/2018 | 05/06/2018 | 13 | 12 | 967 (± 120) | 83,265 (± 35,945) | 93 (± 31) | 40,420 (± 17,449) | 999,184 |
| RJ1.V | 23/02/2018 | 14/06/2018 | 17 | 15 | 2,387 (± 554) | 233,504 (± 49,424) | 100 (± 17) | 37,481 (± 7,933) | 3,502,563 |
| RJ2.B | 15/05/2018 | 28/08/2018 | 16 | 16 | 748 (± 12) | 83,574 (± 19,516) | 112 (± 24) | 60,561 (± 14,142) | 1,337,184 |
| RJ2.V | 08/05/2018 | 31/08/2018 | 18 | 19 | 4,097 (± 1,353) | 453,681 (± 173,901) | 112 (± 19) | 29,422 (± 11,278) | 8,619,954 |
| **3rd Round** | | | | | | | | | |
| RJ1.B | 13/03/2019 | 04/07/2019 | 17 | 12 | 134 (± 0) | 14,483 (± 4,800) | 108 (± 36) | 7,031 (± 2,330) | 173,798 |
| RJ1.V | 01/03/2019 | 12/06/2019 | 16 | 16 | 291 (± 89) | 26,472 (± 10,137) | 91 (± 20) | 4,249 (± 1,627) | 423,558 |
| RJ2.B | 13/06/2019 | 19/09/2019 | 28 | 22 | 71 (± 58) | 5,983 (± 4,681) | 102 (± 30) | 6,729 (± 910) | 204,279 |
| RJ2.V | 30/04/2019 | 13/08/2019 | 16 | 16 | 2,190 (± 533) | 215,596 (± 71,851) | 97 (± 21) | 13,982 (± 4,660) | 3,449,540 |

[a] Number of events (weeks) that release intervention occurred.
[b] Number of release sites per event.
[c] Number of mosquitoes released per event.
[d] Number of mosquitoes released per event per site (i.e. number of mosquitoes in each release tube).
[e] Number of mosquitoes released per event per intervention area ($km^2$).

**Supplementary Table S2. BG-Sentinel traps allocation.** Number of BG-Sentinel traps allocated to each neighborhood section of Ilha do Governador, from the beginning until the end of field monitoring. Revision of schedules and trap numbers was necessary to manage a long-term monitoring activity, and obtain the best possible spatial resolution.

| **Intervention section** | **Number of BG-traps** | | | **Monitoring schedule** | |
|---|---|---|---|---|---|
| | Max. | Min. | Mean (± SD) | Start | End |
| RJ1.V.Bancários | 7 | 3 | 5.9 ± 1.1 | 10/2017 | 03/2020 |
| RJ1.V.Cacuia | 10 | 2 | 6.6 ± 2.9 | 10/2017 | 03/2020 |
| RJ1.V.Cocotá | 4 | 1 | 2.9 ± 0.9 | 10/2017 | 03/2020 |
| RJ1.V.Freguesia | 26 | 1 | 13.1 ± 5 | 10/2017 | 03/2020 |
| RJ1.V.Monero | 9 | 5 | 6.7 ± 2 | 10/2017 | 03/2020 |
| RJ1.V.Pitangueiras | 3 | 2 | 2.4 ± 0.5 | 10/2017 | 03/2020 |
| RJ1.V.Praia da Bandeira | 6 | 2 | 4.1 ± 1.6 | 10/2017 | 03/2020 |
| RJ1.V.Ribeira | 7 | 2 | 4.2 ± 1.5 | 10/2017 | 03/2020 |
| RJ1.V.Tauá | 18 | 8 | 12 ± 4.4 | 10/2017 | 03/2020 |
| RJ1.V.Zumbi | 3 | 1 | 1.8 ± 1 | 10/2017 | 03/2020 |
| RJ1.B | 35 | 1 | 29.2 ± 8.1 | 10/2017 | 03/2020 |
| RJ2.V.Cidade Universitária | 41 | 42 | 41.6 ± 0.5 | 12/2017 | 03/2020 |
| RJ2.V.Galeão | 31 | 1 | 29.3 ± 3 | 12/2017 | 03/2020 |
| RJ2.V.Jardim Carioca | 23 | 1 | 21.4 ± 3.5 | 12/2017 | 03/2020 |
| RJ2.V.Jardim Guanabara | 46 | 3 | 39 ± 12.2 | 12/2017 | 03/2020 |
| RJ2.V.Portuguesa | 18 | 2 | 17.5 ± 2.6 | 12/2017 | 03/2020 |
| RJ2.B | 26 | 15 | 23.1 ± 2.2 | 12/2017 | 03/2020 |
| Tubiacanga | 15 | 3 | 5.4 ± 4.6 | 10/2017 | 03/2020 |

**Supplementary Table S3. Nucleotide sequences of primers and probes.** List of primers and probes used for the molecular diagnostics of *Wolbachia* by qPCR and LAMP, and for *kdr* genotyping.

| PRIMER | NUCLEOTIDE SEQUENCE (5’ → 3’) |
|---|---|
| *Ae. aegypti* RPS17 – qPCR | |
| RPS17S Forward | TCCGTGGTATCTCCATCAAGCT |
| RPS17S Reverse | CACTTCCGGCACGTAGTTGTC |
| RPS17S Probe | **HEX**/CAGGAGGAG/**ZEN**/GAACGTGAGCGCAG/**3IABkFQ** |
| *Ae. aegypti kdr* screening – qPCR | |
| 1016 Forward | CGTGCTAACCGACAAATTGTTTCC |
| 1016 Reverse | GACAAAAGCAAGGCTAAGAAAAGGT |
| 1016 Probe Val+ | **VIC**/CCGCACAGATACTTA/**NFQ** |
| 1016 Probe Ile$^{kdr}$ | **FAM**/CCCGCACAGGTACTTA/**NFQ** |
| 1534 Forward | CGAGACCAACATCTACATGTACCT |
| 1534 Reverse | GATGATGACACCGATGAACAGATTC |
| 1534 Probe Phe+ | **FAM**/ACGACCCGAAGATGA/**NFQ** |
| 1534 probe Cys$^{kdr}$ | **VIC**/AACGACCCGCAGATGA/**NFQ** |
| *Wolbachia* – qPCR | |
| WSPTM2 Forward | CATTGGTGTTGGTGTTGGTG |
| WSPTM2 Reverse | ACACCAGCTTTTACTTGACCAG |
| WSPTM2 Probe | **FAM**/TCCTTTGGA/**ZEN**/ACCCGCTGTGAATGA/**3IAbRQSp** |
| *Wolbachia* – LAMP | |
| FIP | TGTATGCGCCTGCATCAGCTTCGGTTCTTATGGTGCTAA |
| BIP | GCAGAAGCTGGAGTAGCGTTGTGTCATGCCACTTAGATGG |
| F3 | TGATGTAACTCCAGAAGTCA |
| B3 | CTTATTGGACCAACAGGATCG |
| LpF | AGCCTGTCCGGTTGAATT |
| LpB | CAGTCTTGTTATCCCAGTGAGT |
